# Limiting the impact of protein leakage in single-cell proteomics

**DOI:** 10.1101/2024.07.26.605378

**Authors:** Andrew Leduc, Yanxin Xu, Gergana Shipkovenska, Zhixun Dou, Nikolai Slavov

## Abstract

Limiting artifacts during sample preparation can significantly increase data quality in single-cell proteomics experiments. Towards this goal, we characterize the impact of protein leakage by analyzing thousands of primary single cells that were prepared either fresh immediately after dissociation or cryopreserved and prepared at a later date. We directly identify permeabilized cells and use the data to define a signature for protein leakage. We use this signature to build a classifier for identifying damaged cells that performs accurately across cell types and species.

## Main

Recent advances in throughput of single cell proteomics by mass spectrometry have made it possible to quantify thousands of proteins across thousands of single cells^1–4^, including primary cells^3,5,6^. This will facilitate characterizing the influence of protein translation and degradation on shaping the functions of single cells in heterogeneous tissue samples^7^. However, dissociation and long term storage of samples before sample preparation may damage cells^8^ leading to potential bias in single-cell proteomics data.

Indeed, such effects have been observed in single-cell RNA sequencing, where transcripts have been found to leak out of cells with damaged membranes depending upon their localization in or outside of the mitochondria^9^. These cells are usually filtered out computationally based on a heuristic cut off which varies depending on the cell type^9,10^. However, an analogous characterization has not been performed in single-cell proteomics.

Proteins vary significantly in subcellular localizations, physical properties and binding interactions, all of which may substantially affect their leakage propensities. Additionally, proteins are about 10-fold smaller than the mRNA that template them, making them more likely to leak upon membrane damage. For these reasons, we sought to characterize this effect.

We chose to work with primary tissue, mouse tracheal epithelium, to characterize the effect on diverse cell types. After dissociation using an enzyme cocktail as previously reported^11^, half the cells were slowly frozen to -80C in 10% DMSO and 90% FBS and prepared later, and half were prepared fresh immediately for single-cell proteomic analysis. Using nPOP with TMTpro 32-plex multiplexing, we prepared a total of 2,784 single cells, 928 fresh, and 1,856 frozen ones and analyzed them with pSCoPE^5^ using 1,018 cells / day method, quantifying 712 proteins per cell on average. Summary reports of all data can be found in supplementary files 1 (fresh cells) and 2 (frozen cells). Prior to cell isolation, the cells were stained with Sytox green to identify cells with compromised membrane permeability. We recorded the stain intensity of each cell and linked these measurements with downstream single-cell data, Fig 1a. The distribution across all single cells is bimodal, and the cells from the mode at 0 intensity were characterized as intact while the cells from the other mode as permeable, Fig 1a.

**Figure 1.**
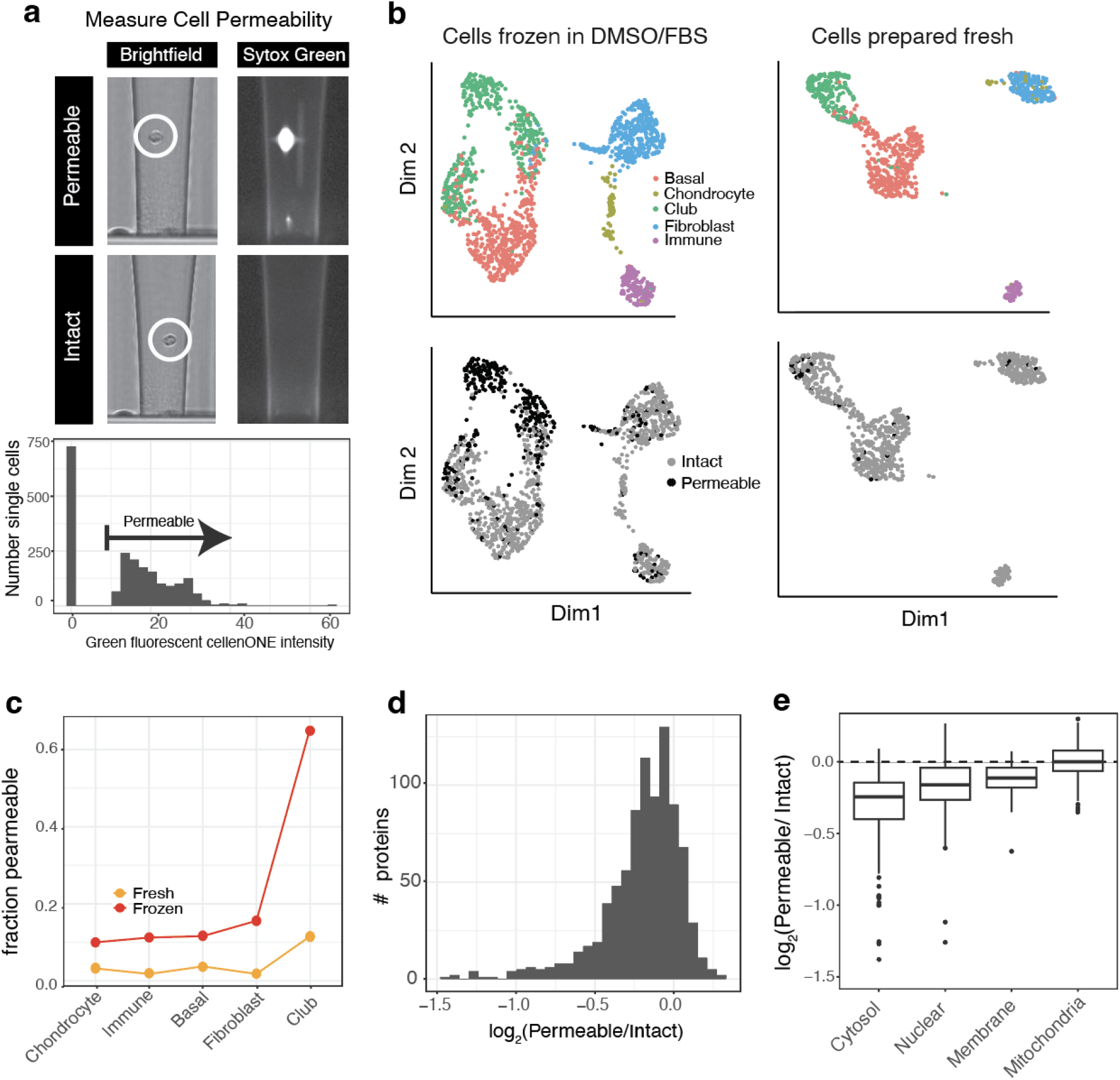
Quantifying protein leakage artifacts. **a**, Images of cells in the CellenONE nozzle taken with brightfield and with the green fluorescent channel. The two cells shown are not clearly distinguishable from the brightfield but one cell is positive for the sytox green fluorescent dye. **b**, The UMAP dimensionality reduction shows co-clustering of cells from fresh and frozen batches that were recorded as permeable via Sytox green. **c**, Percentages of permeable cells for the two sample handling conditions are shown in the dot plot. **d**, Differences in protein fold changes between permeable and intact cells for all proteins. **e**, Differences in protein fold changes between permeable and intact are categorized by subcellular compartment.

Further, the distribution of cell sizes associated with greater than or less than 0 intensity were indistinguishable within a cell type suggesting that cells do not have an absence of signal due to their small size, supplementary Fig 1a. The Sytox green negative cells were 96 % for fresh and 72 % for frozen ones.

We then assigned cell types by using the LIGER^12^ algorithm to perform label transfer from a previously annotated single cell mRNA seq data set^11,12^. The correspondence to the single-cell mRNA data set was lower in frozen samples compared to the fresh. This was observed by reduced agreement in covariance patterns between shared genes^13^, and a reduction in neighborhood similarity score output by LIGER from 0.92 to 0.87, supplementary Fig 2b. We also observed co-clustering of permeabilized cells in the frozen sample, Fig 1b. We did not observe a separate permeable cell cluster in the fresh samples, which may have been due to the low number of permeable cells, Fig 1b. Despite the lower confidence in integration score and co-clustering of permeabilized cells, we were still able to confidently assign cell types in the frozen condition based on the abundance of marker proteins for each cell type, supplementary Fig 1c-f. The proportion of permeable cells was unevenly distributed across cell types, Fig 1c. This may reflect both the increased fragility of the more significantly affected cell types and choices in single cell dissociation.

We next examined which proteins exhibited a significant change between permeable and intact cells, Fig 1d. Most altered proteins were depleted from the permeabilized fraction, indicating that proteins were only lost for the cells characterized as permeable. We also characterized the extent to which the reduction in protein levels was specific to different subcellular compartments. We plotted the difference in protein abundance between different subcellular compartments, Fig 1e. Similar to the trends observed in mRNA sequencing, mitochondrial protein abundance is not significantly different between permeable and intact cells. In contrast, proteins localized to the cytosol have the most significant decrease in abundance between intact and permeable cells. Examining specific proteins reveals metabolic enzymes such as peroxidases and enzymes involved in glycolysis like Gapdh with ∼ 2 fold reduced abundance. A full list of proteins with their fold change differences by cell type can be found in supplementary File 3.

To examine the generalizability of the protein leakage artifact, we estimated the average fold change for each protein between permeable and intact cells between cell types with greater than 5 permeable cells, Fig. 2a. Significant agreement of fold changes across cell types suggests that a similar mechanism of protein leakage is operating across cell types. However, weaker agreement between immune cells and other cell types may reflect a cell type specific component. The similarity in fold changes between cell types led us to explore the utility of a classifier for identifying cells with damaged membranes in data sets where permeability staining was not used. To do this, we trained a classifier on cell permeability status using the abundances of the top 50 most significantly leaking proteins. To validate our model, we first trained and tested it on the same cell type leading to a high success rate of classification on the testing set with an AUC = 0.92, Fig 2b. We then sought to see how well the model could generalize by training it on fibroblasts, basal and immune cells and testing on basal cells. Performance decreased slightly but a significant portion of permeable cells could be identified, AUC = 0.86, Fig 2b. This may be partly the result of cell type dependent leaking that may vary across cell types due to shifts in protein abundance and localization patterns. To facilitate easy usage of the classifier, it was incorporated into the QuantQC R package.

**Figure 2.**
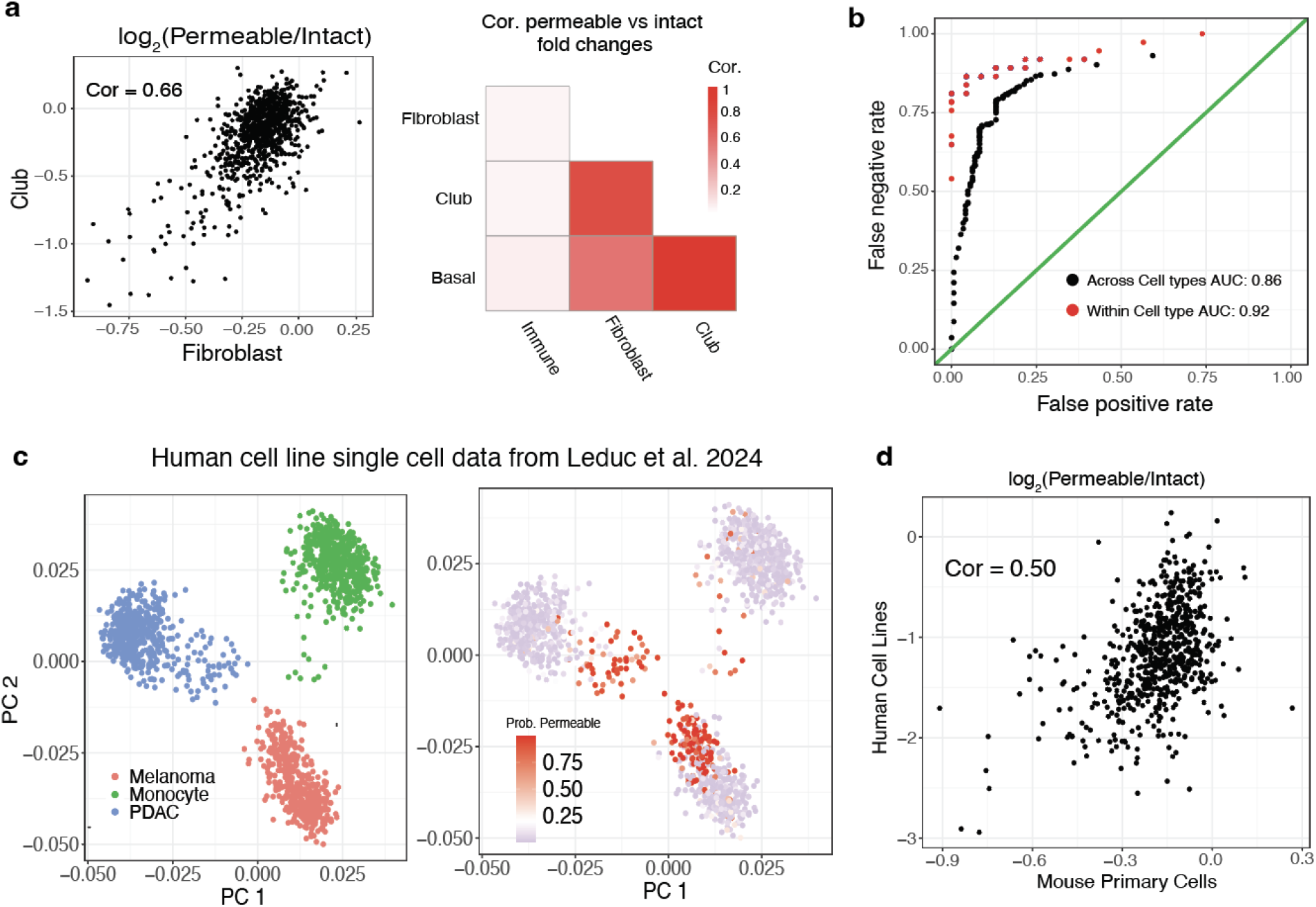
Examining human cell line data. **a**, Log2 average protein fold changes between permeable and intact Club cells and Fibroblasts are plotted against each other and show pearson correlation of 0.66. A heatmap summarizing correlations between fold changes for all cell types is shown. **b**. ROC curve for a classifier trained on permeability status of single cells using protein abundance profiles of the top 75 most significantly leaking proteins. Results for the model trained and tested on the same cell type are in red and trained and tested on different cell types are in black. **c**. PCA projection of single cells from Leduc et al. 2024. Cells are colored by cell type or their permeability score from the classifier. The cells towards the center of the two dimensional space of the first two principal components were enriched for high permeability score. **d**, When comparing the fold changes of cells with probability over 0.2 and under 0.2 to permeable versus intact fold changes from the primary mouse tracheal cells, the fold changes strongly agree, pearson correlation 0.50.

To validate that this phenomenon was generalizable and not just specific to these samples, we next inspected a previously published data set of three human cell lines from our lab ^1^. We trained our classifier for a final time on the entire mouse trachea data set and applied it to the human cells. Upon clustering and low dimensional projection in the principal component space, a population of cells closer to the center of the PCA space from each cell type had high probability of permeabilization, Fig 2c. We then compared the protein abundances between cells with probability greater or less than 0.2 of being permeable as assigned by the classifier. Fold changes between permeable and intact cells from our mouse trachea data set and found strong agreement with the human cell line fold changes, Figure 2d.

Our data demonstrate a substantial impact of protein leakage on single-cell proteomic measurements and a direct resolution based on excluding permeabilized cells from analysis. We also provide protein signatures of leakage and classification tools which may be used to detect and correct for this artifact. While this artifact may be avoidable when performing sample preparation with methods that have fluorescent cell sorting capabilities, this capability is not ubiquitous. Indeed, recent methods have been developed using instruments that lack these capabilities^14^. In such cases, our classification model can help identify and correct for this artifact and thus improve data interpretation. Our results reinforce the importance of incorporating solutions for this problem in the community guidelines and best practices^2^ and point to effective solutions.

## Methods

### Mouse model and handling

All mice experiments were performed in compliance with the Institutional Animal Care and Use Committee at Massachusetts General Hospital. 4-month-old C57BL/6 mice were ordered from the NIA. Mice were euthanized with CO2 followed by cervical dislocation. The mouse used was male. Tissues were harvested post-euthanasia and perfusion with PBS.

### Tissue dissociation and cell suspension generation

Freshly dissected whole trachea preparations were submerged in 500 μl of each enzyme dissociation cocktail for 30 min with gentle rocking at 37 °C. Papain (13.3 U ml−1 or 10 U ml−1) was dissolved in EBSS buffer before mixing with activation buffer consisting of 0.067 mM β-mercaptoethanol, 1.1 mM EDTA and 5.5 mM cystein-HCl in EBSS. Enzyme mix used for the dissociation protocol consisted of 25 μl of 70 kU ml−1 collagenase I, 25 μl of 50 kU ml−1 hyaluronidase, 50 μl of 7.5 kU ml−1 DNase, 120 μl of 2.5 U ml−1 dispase and 400 μl of 40 U ml−1 papain, to a final volume of 5 ml using DMEM. In all cases, single enzyme incubations were done for 30 min at 37 °C with gentle rocking, whereas the enzyme cocktail mix was incubated for 20 min at 37 °C with gentle rocking. Cells were then either taken directly for single cell sample preparation, or frozen down in a cryopreservative buffer of 10% DMSO and 90% FBS.

### Proteomic sample preparation

Samples were prepared using the nPOP sample preparation method for multiplexed single cell proteomics^1^. Briefly, single cells were washed twice from either dissociation buffer or cryopreservation buffer with 1X PBS. Cells were then resuspended at a concentration of 1,000 cells per μL and were incubated on ice and in the dark for 20 minutes with Sytox Green Dead Cell Stain (Thermo Fisher S34860). Cells were then washed one final time to remove dye and resuspended in 1X PBS at a concentration of 300 cells per μL for eventual cell sorting. Cells were then sorted in a volume of 300 pL into 9 μL of 100% DMSO droplets on the surface of a fluorocarbon coated glass slide for cell lysis using the CellenONE cell sorter and liquid handler. As cells were sorted, the fluorescent intensity of the Sytox Green stain was recorded using the CellenONE’s green channel. The remaining cells were pelleted and resuspended in mass spectrometry grade water at a concentration of 1,000 cells per μL and frozen at -80 C. 13.5 μL of digestion buffer of 100 ng/μL trypsin (Promega), 0.025% DDM, and 10 mM HEPES pH 8.5 was added. Samples were incubated for overnight digestion with the aid of the CellenONE’s humidifier and slide cooling for evaporation prevention of droplets.

The next day, samples were labeled with 20 μL of TMTpro 35-plex reagents dissolved in 100% DMSO at a concentration of 8.3 μg/μL. Cells were labeled in sets of 29 as 126C and 127N are reserved for carrier and reference channels, 127C is excluded due to isotopic impurities from the carrier, and we did not have access to the full 35-plex set at the time of the sample preparation. The labeling reaction was then quenched with 20 μL of 1% Hydroxylamine. Samples were pooled using the CellenONE in a 50%/50% solution of Acetonitrile and Water and dispensed into a 384 well PCR plate, dried down in a speed vac and stored at -20C for later injection for LC/MS analysis.

Cells frozen at -80 C in water were then lysed using heat at 90 C for 10 minutes following the mPOP sample preparation ^13^. Cells were then digested over night at 37 C in 100 mM TEAB buffer pH 8.5 and 10ng/μL trypsin. Labels 126C and 127N were used to label carrier and reference samples, respectively. The samples were combined and diluted in 0.1% formic acid to a concentration of 10ng/μL of peptide from the 126C labeled sample for carrier and 0.5 ng/μL from the reference. Samples from the plate were resuspended in 1 uL of the carrier mix for injection.

### LC/MS analysis

Samples were run on an Exploris 480 mass spectrometer with a Vanquish Neo liquid chromatography and autosampler. A 25 cm 75 μm ID Ionoptics column was used for the chromatography column. The gradient ramped from 8-40% buffer B (80% Acetonitrile 20% 0.1 % formic acid) over the course of 28 minutes with a 4 minute wash at 90% buffer B at the end. Mass spectrometry data acquisition of single cells was performed using MaxQuant Live for Prioritized data acquisition with 60k Ms1 and Ms2 resolution, 118 ms maximum injection time and 1e6 maximum AGC ^5^. Briefly, an original inclusion list was generated using a DIA run of the carrier sample which resulted in roughly 10,000 identified precursors. Priority tiers were assigned at three levels, which were equally sized and faceted based on precursor abundance, with highest abundance precursors placed on the highest priority tier. The inclusion list was then refined using two scout runs to identify and remove precursors which had low precursor ion fraction (PIF) below a value of 0.7.

The DIA run for inclusion list generation used a single Ms1 scan and 26 th wide Ms2 scans spanning 400-900 M/Z space with 1 th overlap window to window. The chromatography gradient was the same as specified for the prioritized runs.

### Raw and processed data analysis

Raw data from data dependent and prioritized data acquisition runs were searched by MaxQuant version 2.4.3.0 against a protein sequence database including entries from the appropriate human SwissProt database (downloaded July 30, 2018) containing 20,386 proteins and known contaminants such as human keratins and common lab contaminants. The modifications for the TMTpro 35plex tags can be found in the supplemental data of the nPOP protocol ^1^. Results were filtered at 1% FDR. The DIA run for inclusion list generation was searched using the DIA-NN software with TMT specified as a fixed modification.

Downstream data analysis was performed in R. Single cell data was processed to obtain a protein X single cell matrix of log2 relative protein fold changes using the QuantQC package ^1^. Cell types were then assigned by integrating of single cell data with pre-annotated mRNA sequencing data from the same tissue type and dissociation procedure ^11^. The LIGER algorithm was used to project the cells into the same high dimensional space. Clustering was performed and protein single cells were assigned the identity of the predominant annotated mRNA single cells from the given cluster. Fold changes were then calculated between permeable and intact cells by taking the average fold change of permeable and intact cells within a cell type for each protein and subtracting the two vectors.

### Cell permeability classifier

The model used to classify permeable vs intact single cells XGboost was trained on the top 75 most significantly leaking proteins (Supplemental file 3) across single cells. Data was z-scored within each protein (across single cells) prior to training the model.

Missing data was left as NA. For the within cell type comparison, the 420 club cells from the Frozen data set were randomly split into train and test sets of 80 and 20 percent of the data, respectively. For the across cell type comparison, the model was trained on all cells except for club cells and then tested on all club cells.

## Data and Code Availability

The analysis can be reproduced by using resources from https://scp.slavovlab.net/Leduc_et_al_2024 and following instructions found at https://github.com/SlavovLab/CellPermeability. The classification tool has been incorporated into the QuantQC package, https://github.com/SlavovLab/QuantQC via the function FindPermeableCells.

The raw and searched MS data have been deposited in accordance with community guidelines^2^ and can be found at MassIVE repository MSV000094790 with password nPOP1234.

The processed data can be found at Figshare:

https://figshare.com/articles/figure/Figure_1/25824157/2

And at scp.slavovlab.net:

https://drive.google.com/drive/folders/1IW3An_b7vKpfDbanjT2XqSt1mz0N6VUX

Supplemental files 1-3 can be found at the following google drive link:

https://drive.google.com/drive/folders/1CKhoOD7_VvojQzt9AYKFH3neoTvSzdf9?usp=sharing

## Author contributions

Experimental design: A.L and N.S. Sample preparation: A.L., Y.X., G.S., and Z.D. LC–MS/MS: A.L. Data analysis and writing: A.L and N.S. All authors approved the final manuscript.

## Acknowledgments

We thank Sarah Sipe for help with MS data acquisition at PTI and members of the Slavov laboratory for thoughtful discussions. The work was funded by a Bits to Bytes award from MLSC to N.S., an NIGMS award R01GM144967 to N.S., and a MIRA award from the NIGMS of the NIH (R35GM148218) to N.S, and a UH3CA268117 award from NIH to N.S and Z.D.

---

**Supplemental figure 1.**
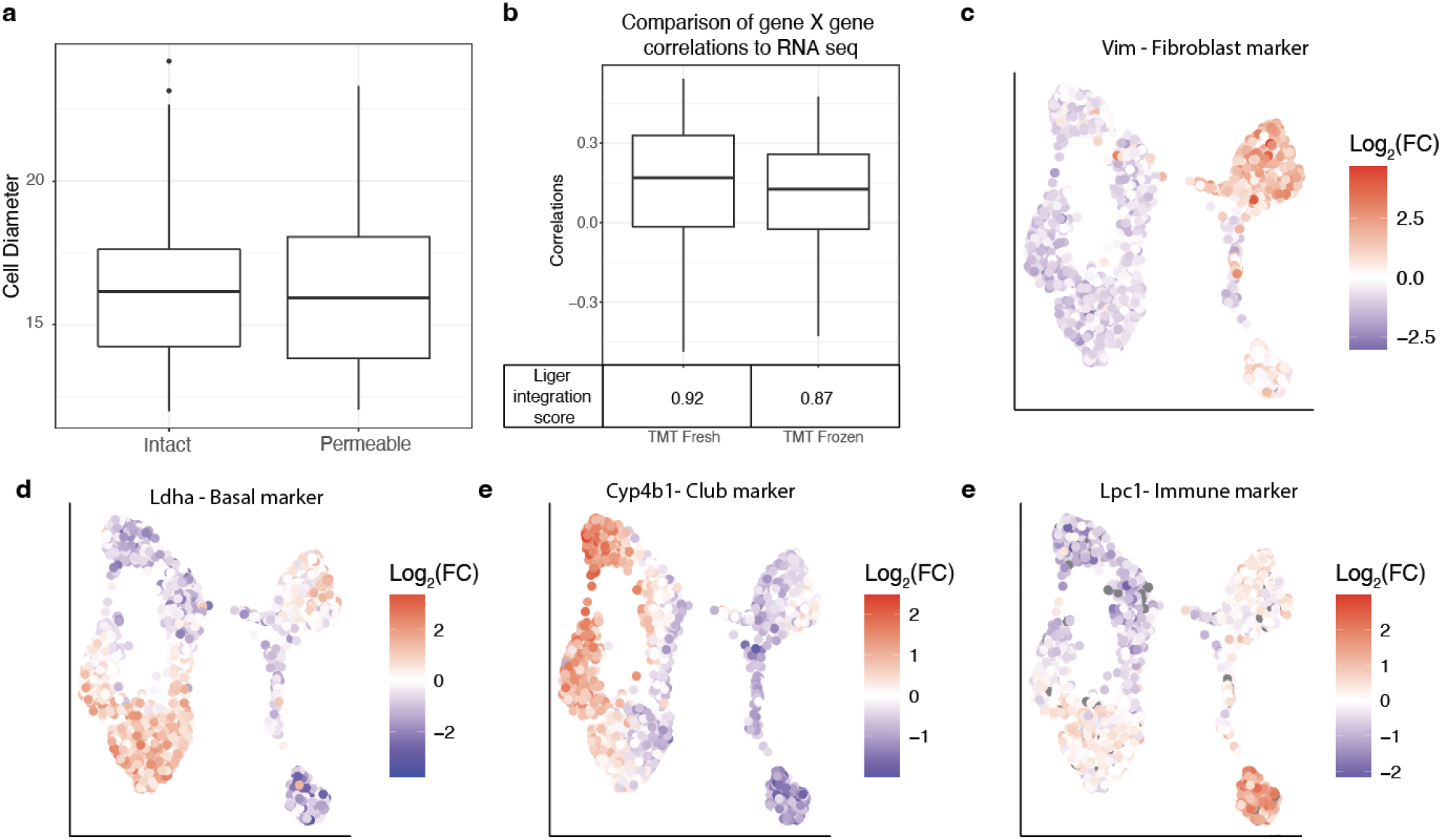
Assigning permeability and cell types. **a**, CellenONE diameters for intact and permeable cells. **b**, Protein-protein and transcript-transcript correlation matrices were computed for intersected genes. The correlation vectors for like genes were correlated. Distributions for this comparison in the fresh and frozen data sets are plotted. The LIGER integration score was .05 higher for the fresh cell data set. **c-f**, UMAP dimensionality reduction plots with the abundance of marker proteins for basal, club, immune and fibroblast cell populations in the frozen sample.

